# Improved genome assembly and annotation of the soybean aphid (*Aphis glycines* Matsumura)

**DOI:** 10.1101/781617

**Authors:** Thomas C. Mathers

## Abstract

Aphids are an economically important insect group due to their role as plant disease vectors. Despite this economic impact, genomic resources have only been generated for a small number of aphid species. The soybean aphid (*Aphis glycines* Matsumura) was the third aphid species to have its genome sequenced and the first to use long-read sequence data. However, version 1 of the soybean aphid genome assembly has low contiguity (contig N50 = 57 KB, scaffold N50 = 174 KB), poor representation of conserved genes and the presence of genomic scaffolds likely derived from parasitoid wasp contamination. Here, I use recently developed methods to reassemble the soybean aphid genome. The version 2 genome assembly is highly contiguous, containing half of the genome in only 40 scaffolds (contig N50 = 2.00 Mb, scaffold N50 = 2.51 Mb) and contains 11% more conserved single copy arthropod genes than version 1. To demonstrate the utility this improved assembly, I identify a region of conserved synteny between aphids and *Drosophila* containing members of the *Osiris* gene family that was split over multiple scaffolds in the original assembly. The improved genome assembly and annotation of *A. glycines* demonstrates the benefit of applying new methods to old data sets and will provide a useful resource for future comparative genome analysis of aphids.

## Introduction

Aphids are an economically important insect group due to their role as plant disease vectors (Van Emden and Harrington 2017). They are also important models used to study plant-insect interactions (Hogenhout and Bos 2011; Ferry et al. 2004), speciation genomics (Pecoud and Simon 2010; Peccoud et al. 2009; Hawthorne and Via 2001) and sex chromosome evolution (Jaquiéry et al. 2018, 2013). Despite their importance, only a small number of the approximately 5,000 described aphid species have had their genomes sequenced (IAGC 2010; Nicholson et al. 2015; Wenger et al. 2017; Mathers et al. 2017; Chen et al. 2019; Jiang et al. 2019; Quan et al. 2019; Thorpe et al. 2018), limiting genomic insights into their diversity and evolution. Furthermore, although highly contiguous assemblies have recently been published for two aphid species (Jiang et al. 2019; Chen et al. 2019), the majority of publicly available aphid genomes were sequenced using second-generation short-read sequencing technology, resulting in fragmented assemblies that contain thousands of genomic scaffolds. Although these assemblies may be accurate at the gene-level, and have facilitated many important discoveries, they likely underrepresent repetitive genome content (Treangen and Salzberg 2012; Sedlazeck et al. 2018) and may be unsuitable for analyses such as the detection of large-scale structural variants (Chaisson et al. 2015; Chakraborty et al. 2017) and genome-wide synteny analysis (Liu et al. 2018). To gain a fuller understanding of aphid evolution and adaptation many more high-quality genomes are required. This will primarily be achieved through new genome sequencing projects. However, as improved informatic approaches are developed, reuse of existing data sets will also make a useful contribution to improving genomic resources available for aphids and other taxa.

The soybean aphid (*Aphis glycines* Matsumura) is an important introduced crop pest in North America and was the third aphid species to have its genome sequenced and the first to use long-read sequence data (Wenger et al. 2017; Tilmon et al. 2011). Version 1 of the *A. glycines* genome assembly (herein referred to as A_gly_v1) was assembled using genomic libraries prepared from wild-caught samples identified as biotype 1 or 4 (n=21) from across the USA, and from a single library derived from a lab strain of biotype 4 (Wenger et al. 2017). The wild-caught samples were sequenced using short-read technology (Illunmina MiSeq, 300 bp PE, ∼147 x genome coverage) and the biotype 4 lab colony was sequenced using the Pacific Biosciences (PacBio) single molecule real time sequencing (SMRT) platform to generate long-read data (∼25x genome coverage). Wenger et. al. (2017) combined all the short-read sequence data from both biotypes to generate an initial *de novo* assembly that was subsequently scaffolded using the PacBio long reads. This produced a fragmented genome assembly containing 8,397 scaffolds totalling 301 Mb of sequence with a scaffold N50 of 174 Kb (contig N50 = 57 Kb).

Genome assembly algorithms are a source of constant innovation and improvement (Sedlazeck et al. 2018). This is particularly true in the field of long-read genome assembly and in the integration of short- and long-read data (hybrid genome assembly). However, despite the public deposition of data, genome assemblies of non-model organisms are rarely revisited. Here, I leverage recently developed methods to reassemble the soybean aphid genome using the original sequence data. The version 2 genome assembly is highly contiguous, containing half of the genome in only 40 scaffolds (contig N50 = 2.00 Mb, scaffold N50 = 2.51 Mb), and has improved accuracy at the gene-level, with the representation of conserved single copy arthropod genes (n=1,066) increased by 11% compared to version 1 (983 vs 888). To demonstrate the utility of the updated genome assembly, I investigate synteny of the insect-specific gene family *Osiris* (Shah et al. 2012) between *Drosophila melanogaster* and *A. glycines*. The updated assembly of *A. glycines* resolves the complete *Osiris* gene cluster and reveals conserved synteny between aphids and flies over approximately 250 million years. The updated genome assembly and annotation of *A. glycines* will provide solid foundation to understand the biology of *A. glycines* and other aphid species.

## Results and Discussion

### Assessment of *Aphis glycines* v1

I assessed the quality of A_gly_v1 and a selection of published aphid genome assemblies by searching for conserved single copy genes using BUSCO (Simão et al. 2015; Waterhouse et al. 2018) with the Arthropoda gene set (n=1066). A_gly_v1 contains full length copies of 93.9% of arthropod BUSCOs (**Figure 1a**), indicating a high level of genome completeness. However, compared to other aphid genome assemblies, A_gly_v1 has more than twice as many duplicated BUSCO genes (10.6% vs. 2.3 – 4.7%). This unusual result could be a genuine biological phenomenon or an artefact of the assembly process caused by fragmentation, un-collapsed heterozygosity (separately assembled alleles) or contamination.

**Figure 1:**
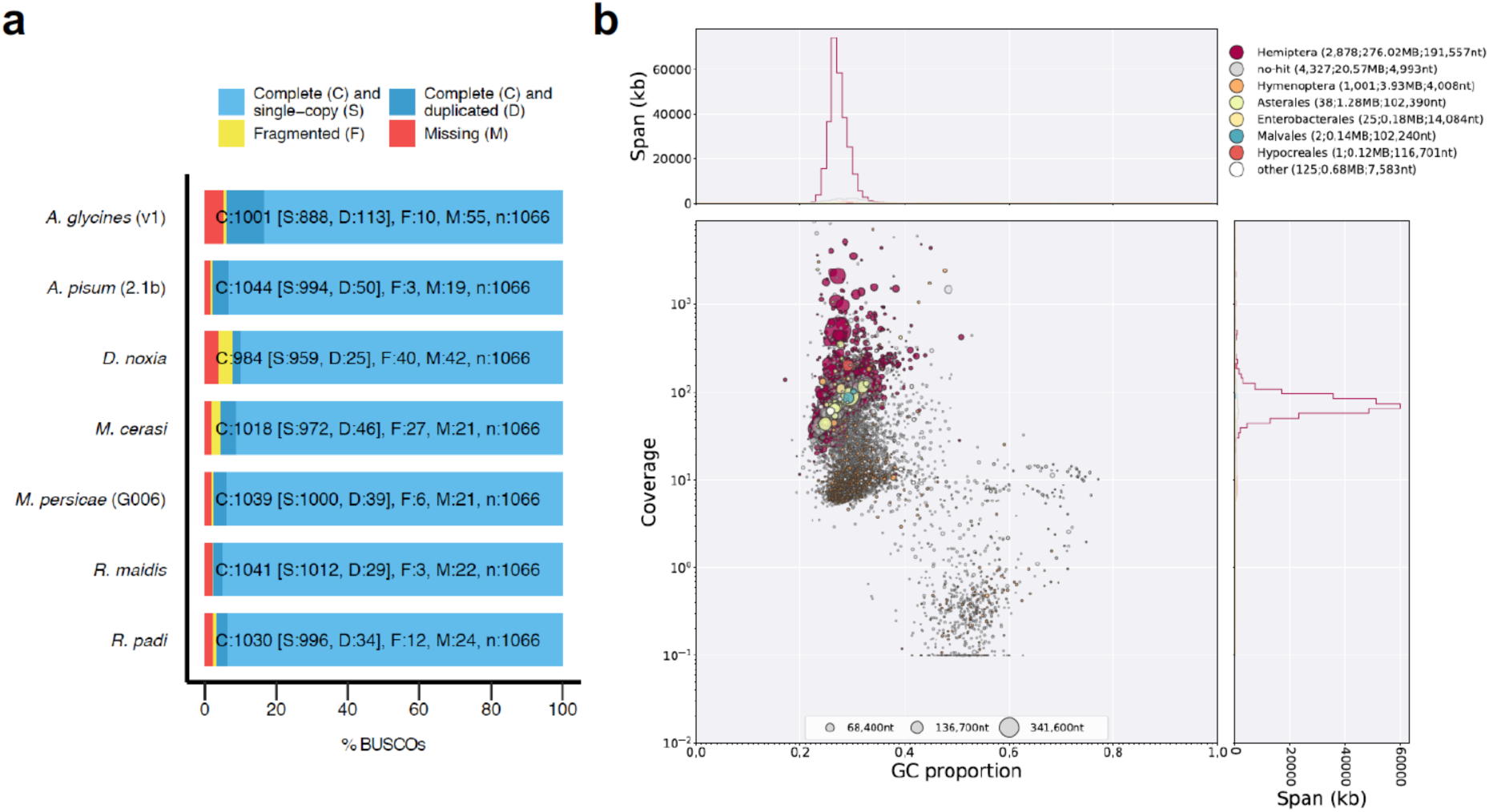
Assessment of the *Aphis glycines* v1 (A_gly_v1) genome assembly. (**a**) BUSCO analysis of published aphid genome assemblies using the Arthropoda gene set of 1,066 conserved single copy genes. Bars show the proportions of genes found in each assembly as a percentage of the total gene set. (**b**) Taxon-annotated GC content-coverage plot of A_gly_v1. Each circle represents a scaffold in the assembly, scaled by length, and coloured by order-level NCBI taxonomy assigned by BlobTools. The X axis corresponds to the average GC content of each scaffold and the Y axis corresponds to the average coverage based on alignment of pooled *A. glycines* Illumina MiSeq short-read libraries from Wenger *et. al.* (2017). Marginal histograms show cumulative genome content (in Kb) for bins of coverage (Y axis) and GC content (X axis).

Given that few (0.9%) BUSCO genes are fragmented in A_gly_v1, and that *A. glycines* has low genetic diversity across its introduced range in North America (Wenger et al. 2017; Orantes et al. 2012), I focussed on contamination as the likely source of elevated duplication levels. To identify contamination in A_gly_v1 I generated a taxon-annotated GC content-coverage plot (known as a “blob plot”) of all A_gly_v1 scaffolds with BlobTools (McLean et al. 2018). This revealed two distinct “blobs”, and an additional group of scaffolds with low coverage and high GC content, indicating the presence of contamination (**Supplementary Table 1**; **Figure 1b**). Scaffolds in the primary “blob” account for the majority of A_gly_v1 sequence and are mostly assigned to Hemiptera as expected. Scaffolds in the secondary “blob” are assigned to Hymenoptera and have lower average coverage (118x vs 28x) and higher average GC content (32.5% vs 27.8%) than those assigned to Hemiptera, indicating they are derived from different genomes (Kumar et al. 2013). The likely explanation for this is that some sequence libraries used in the original assembly were derived from wild-caught aphids infected with parasitoid wasp larvae. In total, 1,001 out of 8,397 A_gly_v1 scaffolds (totalling 3.93 Mb of sequence) are assigned to Hymenoptera. This likely represents an underestimation of the hymenopteran content in A_gly_v1 as there are many scaffolds with unannotated taxonomy also clustering with the Hymenoptera scaffolds due to a lack of sequenced aphid parasitoid wasp genomes. Consistent with this, blast hit identities for Hymenoptera scaffolds are significantly lower than for Hemiptera scaffolds (Mann-Whitney U Test: p < 2.2×10^−16^, U = 1552300000; **Supplementary Figure 1**). Nonetheless, inspection of the A_gly_v1 official gene set (v1.0) reveals that scaffolds assigned to Hymenoptera contain 806 genes previously thought to be derived from *A. glycines*. These scaffolds also account for 68 out of 113 of the duplicated BUSCO genes. This is a serious problem as contaminated genome assemblies have the potential to significantly affect downstream comparative analysis (e.g. Koutsovoulos et al. 2016).

### Reassembly of *A. glycines* biotype 4

A_gly_v1 was assembled from data derived from wild-caught aphids (short-read data, ∼147x coverage) and from a lab-reared biotype 4 colony (PacBio long-read data, ∼25x coverage) (Wenger et al. 2017). I reasoned that the lab-reared colony was unlikely to be contaminated with parasitoid wasps and could be used to generate a clean *A. glycines* genome assembly. I therefore took advantage of improvements to the Canu genome assembler (Koren et al. 2017) that enable assembly of low coverage long-read datasets, to generate an initial *de novo* assembly of the *A. glycines* PacBio dataset. This resulted in 1,967 contigs, totalling 301 Mb of sequence, with an N50 of 409 Kb. Contigs from this assembly formed a single blob around the expected GC content, with no evidence of parasitoid wasp contamination (**Supplementary Figure 2**).

Although the Canu assembly is approximately 7 times more contiguous than A_gly_v1 (409 Kb vs. 57 Kb; **Figure 2a**), further gains may still be possible using alternative assembly strategies. It has been shown that combining PacBio-only assemblies with hybrid assemblies that use both long- and short-read data can dramatically improve contiguity due to the complementary assembly of different genome regions (Chakraborty et al. 2016). To produce a hybrid assembly of *A. gylcines* biotype 4, I first identified contamination free *A. glycines* short-read libraries sequenced by Wenger *et. al.* (2017) to be used to create an accurate de Bruijn graph-based short-read assembly to act as seed for a PacBio hybrid genome assembly. Excluding contamination before assembly is preferable to post-assembly filtering in this instance as the parasitoid wasp and aphid have similar GC content (**Figure 1b**), making it difficult to distinguish between target species contigs and contamination. To identify libraries that contain high levels of contamination, I mapped the Illumina libraries derived from wild-caught biotype 4 aphids (n=13) to the Canu assembly and set aside libraries with low mapping efficiency (< 75% of reads mapped). Eleven biotype 4 Illumina libraries passed this filtering step (**Supplementary Figure 3**) and were assembled with DISCOVAR *de novo* (Weisenfeld et al. 2014). This short-read assembly was then used as an input to DBG2OLC (Ye et al. 2016) to generate a hybrid assembly with the PacBio dataset. The DBG2OLC assembly contained 824 contigs totalling 284 Mb of sequence with a contig N50 of 703 Kb. Next, I used quickmerge (Chakraborty et al. 2016) to combine the Canu and DBG2OLC assemblies. This further increased contiguity, producing a merged assembly containing 1,068 contigs totalling 306 Mb of sequence with a contig N50 of 2.06 Mb, a 35-fold increase in contig-level N50 compared to A_gly_V1 (**Figure 2a**). This assembly was then subjected to two rounds of polishing with Pilon (Walker et al. 2014) followed by filtering to remove contigs derived from mitochondrial DNA, and *Buchnera* and *Wolbachachia* endosymbionts (**Supplementary Figure 4**). Finally, the polished and filtered assembly was scaffolded using *A. glycines* RNA-seq data (Bansal et al. 2014) with P_RNA_scaffolder (Zhu et al. 2018) to produce version 2 of the *A. glycines* genome assembly (A_gly_v2).

**Figure 2:**
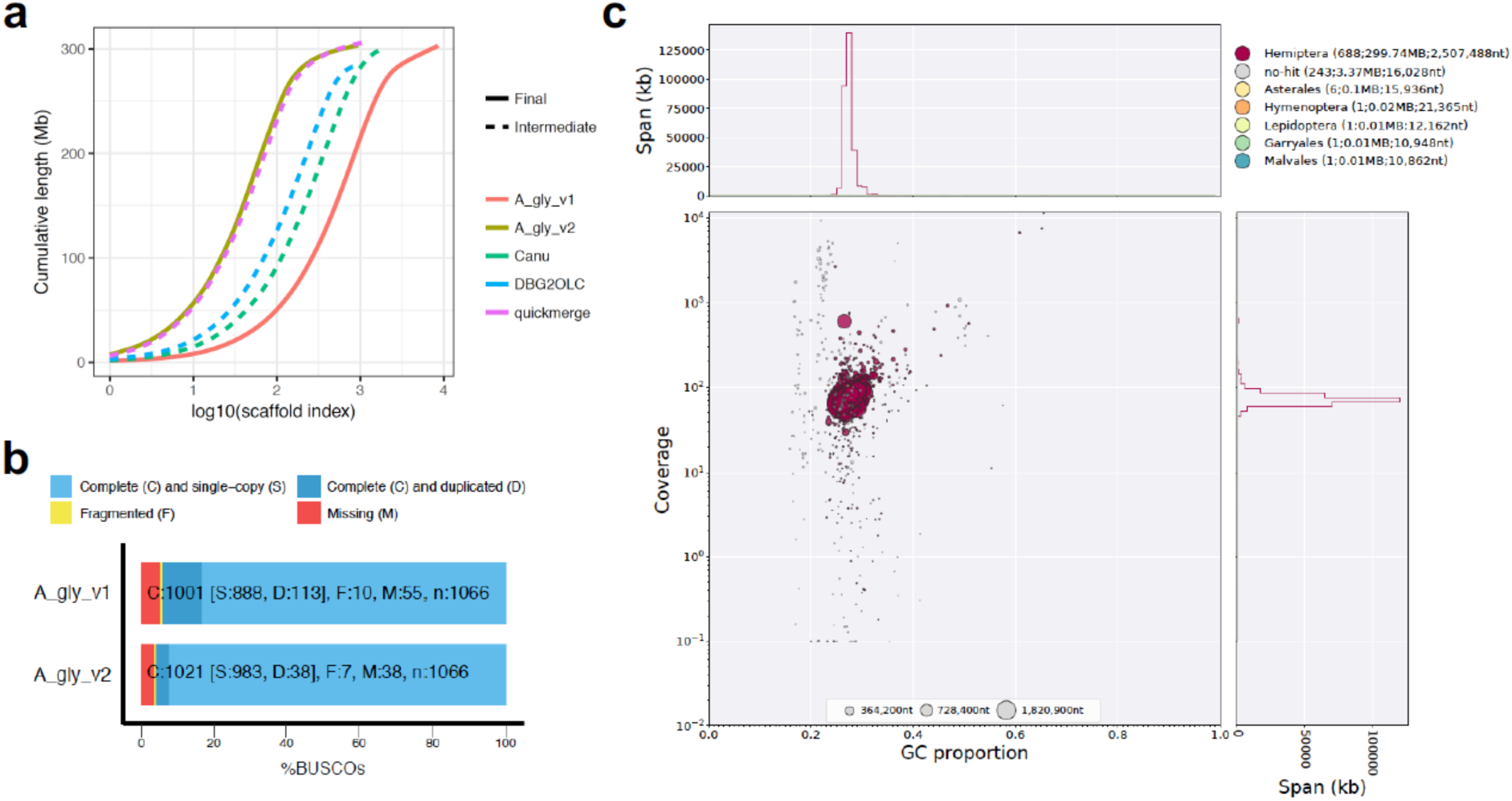
The updated assembly of *A. glycines* (A_gly_v2) is contiguous, complete, and free from contamination. (**a**) Cumulative scaffold (or contig) length for the original assembly of *A. glycines* (A_gly_v1), the PacBio only assembly (Canu), the hybrid assembly (DBG2OLC), the merged PacBio only and hybrid assembly (quickmerge) and the final updated assembly that includes scaffolding with RNA-seq data and removal of contaminants (A_gly_v2). Summary statistics for each assembly are given in **Table 1**. (**b**) BUSCO analysis of A_gly_v1 and A_gly_v2 using the Arthropoda gene set (n=1,066). (**c**) Taxon-annotated GC content-coverage plot of A_gly_v2.

A_gly_v2 is contiguous, free from obvious contamination and is highly complete. Half of the genome is contained in only 40 scaffolds (941 scaffolds in total, longest scaffold = 7.28 Mb) and the scaffold N50 is increased by 1,342% (2.51 Mb vs 0.17 Mb) compared to A_gly_v1 (**Table 1**;**Figure 2a**). After exclusion of parasitoid wasp scaffolds from A_gly_v1, A_gly_v2 contains 4 Mb more sequence than A_gly_v1 (303 Mb vs. 299 Mb), and is close to the predicted genome size of 317 Mb based on flow cytometry (Wenger et al. 2017). Furthermore, A_gly_v2 contains 95 more single copy and complete BUSCO genes than A_gly_v1 (**Figure 2b**; 983 vs. 888) and is free from obvious contamination of non-target species (**Figure 2c**). This improvement in gene content is the result of a reduction in the number of fragmented or missing genes (45 vs. 65) and greatly reduced duplication levels (38 vs. 113). Using BRAKER2 (Hoff et al. 2015; github.com/Gaius-Augustus/BRAKER) with RNA-seq evidence from Bansal *et. al.* (2014), I annotated 19,750 protein coding genes in A_gly_v2 to generate a new gene release to accompany the updated genome assembly.

**Table 1:** Assembly and annotation statistics for the original *A. glycines* genome assembly (A_gly_v1) and alternative assemblies of *A. glycines* generated in this study (see main text).

| Assembly | A_gly_v1 | Canu | DBG2OLC | quickmerge | A_gly_V2 |
| --- | --- | --- | --- | --- | --- |
| Base pairs (Mb) | 302.92 | 301.09 | 282.98 | 305.95 | 303.15 |
| % Ns | 0.14 | 0.00 | 0.00 | 0.00 | 0.00 |
| Number of contigs | 24,335* | 1,967 | 863 | 1,080 | 1,024* |
| Contig N50 (Kb) | 57 | 409 | 703 | 2,064 | 2,000 |
| Number of scaffolds | 8,397 | NA | NA | NA | 941 |
| Scaffold N50 (Kb) | 174 | NA | NA | NA | 2,507 |
| Longest scaffold (Mb)** | 1.37 | 1.78 | 3.10 | 6.26 | 7.28 |
| Protein coding genes | 19,182 | NA | NA | NA | 19,750 |
| Transcripts | 19,182 | NA | NA | NA | 21,647 |

\*Scaffolds split on runs of 1 or more Ns
\*\*Longest contig shown if no scaffolding has been carried out.

### Resolution of the aphid *Osiris* gene cluster

*Osiris* is a large insect-specific gene family that has retained high levels of synteny during insect evolution (Shah et al. 2012; Smith et al. 2018), making it ideal for assessing genome assembly quality. I annotated *Osiris* gene family members in A_gly_v1 and A_gly_v2 based on genome-wide synteny analysis with *Drosophila melanogaster*, blast searches with *D. melanogaster Osiris* genes and phylogenetic analysis (**Supplementary Figure 5**; **Supplementary Table 2**). In total, I identified 29 *Osiris* genes in A_gly_v2. Of these, 19 are located on a single 5 Mb scaffold (scaffold_7). In comparison, scaffold_7 is broken up into 19 parts in A_gly_v1, with the *Osiris* gene cluster split across 2 scaffolds (**Figure 3a and b**). Synteny with *D. melanogaster* on A_gly_v2 scaffold_7 is limited to the region containing *Osiris* genes (**Supplementary Figure 6**) and, given that A_gly_v2 scaffold_7 extends up- and down-stream several Mb, it appears that the entire *A. glycines Osiris* gene cluster has been resolved (**Figure 3a and b**). The increased contiguity of A_gly_v2 reveals rearrangements in gene order at the extremity of the *Osiris* gene cluster, where *Osiris* 2 and 24 have been shuffled to the 3’ end in A_gly_v2 relative to *D. melanogaster* (**Figure 3b**). This rearrangement was previously hidden by genome assembly fragmentation in A_gly_v1, and in the original pea aphid genome assembly, where *Osiris* gene order has also been investigated (Shah et al. 2012).

**Figure 3:**
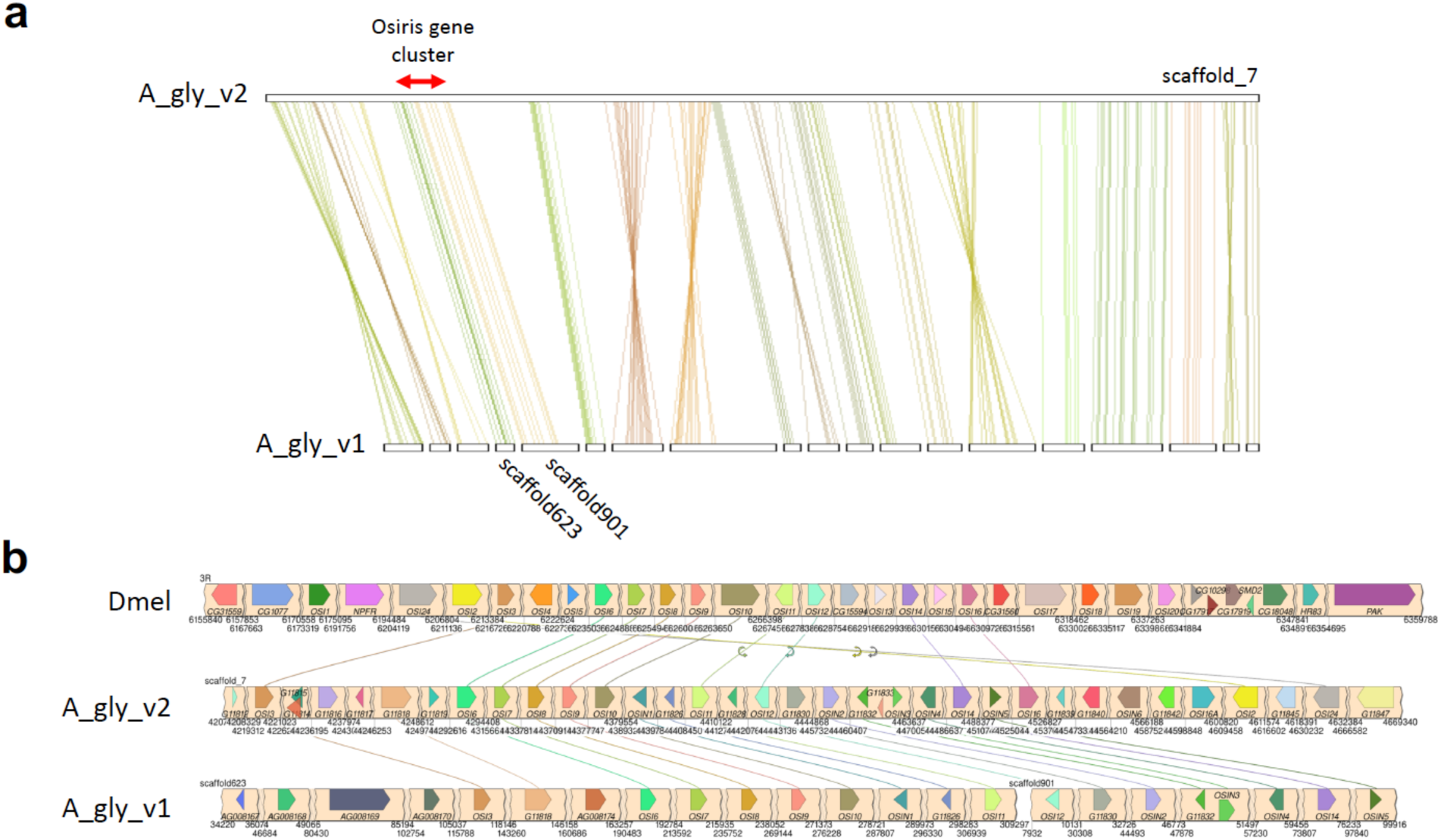
The *A. glycines Osiris* gene cluster is resolved in A_gly_v2 and shares synteny with *D. melanogaster*. (**a**) MCScanX gene-level colinearity between A_gly_v2 scaffold_7 and multiple scaffolds from A_gly_v1. The position of the *Osiris* gene cluster on A_gly_v2 scaffold_7 is indicated by a red arrow. This region is split across two scaffolds in A_gly_v1. (**b**) Schematic showing synteny between *D. melanogaster*, A_gly_v2 and A_gly_v1 for the *Osiris* gene cluster. *A. glycines Osiris* genes are named according to the annotation in **Supplementary Table 2**. Links are shown between one to one *Osiris* orthologs.

Genome-wide synteny between *A. glycines* and *D. melanogaster* is restricted to two regions, one containing the *Osiris* gene cluster, and another containing a cluster of GMC oxidoreductase genes (**Supplementary Figure 6**), further highlighting the extraordinary conservation of *Osiris* gene order within insects over hundreds of millions of years (Shah et al. 2012). Factors that may have selected for the long term conservation of *Osiris* gene order remain elusive (Shah et al. 2012; Smith et al. 2018). However, several lines of evidence indicate *Osiris* genes may be important in the biology of aphids and other plant-feeding insects. *Osiris 7* has been identified in the saliva of the wheat-feeding aphid *Diuraphis noxia*, indicating it could be involved in insect-plant interactions (Nicholson et al. 2012), and *Osiris* 5 is associated recurrent adaptation to toxic fruit in *Drosophila* (Yassin et al. 2016). In *A. glycines*, phylogenetic analysis shows expansion of genes related to *Osiris 5* and *Osiris 16*, with six copies present (**Supplementary Figure 5**). This expansion is likely a conserved feature of aphid genomes as six copies of *Osiris 5 / 16* are also found in *A. pisum* (Shah et al. 2012). The role of *Osiris* gene family members in aphid-plant interactions warrants further study.

## Conclusions

Reassembly of the soybean aphid genome highlights the potential to gain new insights by revisiting old datasets and applying new analysis approaches. Through curation and reanalysis of existing data I have achieved a large increase in genome contiguity and completeness relative to A_gly_V1. This new genome assembly of *A. glycines* adds to the small but growing collection of high-quality aphid genome assemblies and will provide a solid foundation for future studies of *A. glycines* biology, and for comparative genomic analysis with other aphid species.

## Methods

### Assessment of *A. glycines* v1 and other aphid genomes

I assessed gene-level completeness of *A. glycines* v1 (A_gly_v1) and other published aphid genome assemblies using Benchmarking Universal Single-Copy Orthologs (BUSCO) v3.0 (Waterhouse et al. 2018; Simão et al. 2015) with the Arthropoda gene set (n = 1,066) using default settings. I obtained the publicly available genome assemblies of *A. glycines* v1 (A_gly_v1) (Wenger et al. 2017), *Acyrthosiphon pisum* (LSR1) v2.1b (IAGC 2010), *Diuraphis noxia* (Nicholson et al. 2015), *Myzus cerasi* (Thorpe et al. 2018), *Myzus persicae* (G006) v1.1 (Mathers et al. 2017) and *Rhopalosiphum padi* (Thorpe et al. 2018) from AphidBase (bipaa.genouest.org/is/aphidbase/). I also included the recently published high quality assembly of *Rhopalosiphum maidis* (Chen et al. 2019).

To check for contamination in A_gly_v1, I generated a taxon-annotated GC content-coverage plot using BlobTools v0.9.19 (Laetsch and Blaxter 2017; Kumar et al. 2013). Each scaffold in A_gly_v1 was annotated with taxonomy information based on blastn v2.2.31 (Camacho et al. 2009) searches against the NCBI nucleotide database (nt, downloaded 13/10/2017) with the options “-outfmt ‘6 qseqid staxids bitscore std sscinames sskingdoms stitle’ -culling_limit 5 - evalue 1e-25”. To calculate average coverage per scaffold, I mapped Illumina MiSeq paired-end sequence data derived from wild-caught *A. glycines* biotype 4 samples (13 libraries) from Wenger et. al. (2017) to A_gly_v1 using BWA-MEM v0.7.7 (Li 2013) with default settings. The resulting BAM file was sorted with SAMtools v1.3 (Li et al. 2009) and passed to BlobTools along with the table of blastn results.

### Reassembly of *A. glycines* biotype 4

To create an initial contamination free assembly of *A. glycines*, I *de novo* assembled the PacBio long-read sequence data from Wenger et. al. (2017) with Canu v1.6 (Koren et al. 2017) using recommended settings for low coverage sequence data: “genomeSize=317m corMinCoverage=0 correctedErrorRate=0.105 gnuplotTested=true ovsMethod=sequential - pacbio-raw”. These data were derived from a lab reared colony of biotype 4 which was unlikely to be contaminated by parasitoid wasp larvae. The Canu assembly was checked for contamination by creating a GC-content coverage plot using KAT sect from the K-mer analysis toolkit (KAT) (Mapleson et al. 2017). For this analysis, all biotype 4 Illumina MiSeq libraries from Wenger et. al. (2017) were used to calculate median K-mer coverage per scaffold.

To generate a hybrid PacBio-Illumina assembly *of A. glycines* biotype 4 I first identified libraries that were likely the source of contamination in A_gly_v1 using the contamination-free Canu assembly. I mapped each biotype 4 Illumina MiSeq library to the Canu assembly with NextGenMap v0.5.5 (Sedlazeck et al. 2013) using default settings and discarded libraries where less than 75% of reads aligned as valid pairs. The retained libraries were then trimmed for adapter sequence with Trim_galore! v0.4.5 (“--quality 0 --paired --length 150”) (Krueger 2015), concatenated, and assembled with DISCOVAR *de novo* (Weisenfeld et al. 2014) with default settings. The discovar assembly was then used as the input for hybrid genome assembly with DBG2OLC (Ye et al. 2016). DBG2OLC was run using all the PacBio data from Wenger et. al. (2017) and with the following settings: “k 17 KmerCovTh 2 MinOverlap 20 AdaptiveTh 0.002 LD1 0 MinLen 200”. A consensus of the resulting backbone assembly was then generated using BLASR (Chaisson and Tesler 2012) and PBDagCon (github.com/PacificBiosciences/pbdagcon) as per recommendations in Chakraborty *et. al.* (2016). The DBG2OLC consensus assembly was then checked for contamination using a GC-content coverage plot with KAT sect as for the Canu assembly.

To create the final *A. glycines* biotype 4 contig assembly I merged the Canu and DBG2OLC hybrid assemblies with quickmerge v0.2 (Chakraborty et al. 2016). I used the Canu assembly as the “query” and the DBG2OLC assembly as the “reference” so that the majority of content in the merged assembly would be derived from the Canu assembly (see quickmerge documentation). The length cut-off for anchor contigs (“-l”) was set to the N50 of the Canu assembly (409,248) and the minimum alignment length (“-lm”) was set to 10 kb as per recommendations in the quickmerge documentation, all other settings were left as default. The merged assembly was then polished with the retained biotype 4 Illumina MiSeq libraries using two rounds of Pilon v1.22 (Walker et al. 2014).

### Assembly filtering and detection of endosymbionts

The polished quickmerge assembly was checked for contamination using two runs of BlobTools, one with per contig coverage calculated using all the available biotype 4 Illumina MiSeq libraries mapped to the assembly with BWA-MEM v0.7.7 (Li 2013), and a second run using the PacBio long-reads mapped to the assembly with minimap2 r672 (Li 2018). blastn searches for the quickmerge assembly BlobTools runs were performed as per A_gly_v1 against the NCBI nt database. The BlobTools analyses revealed the presence of contigs assigned to *Buchnera* and *Wolbachia* endosymbionts (**Supplementary Table 4**), these were filtered from the final assembly. Manual inspection of the BlobPlot also revealed a 1.40 Mb contig assigned to Rickettsiales that had GC content and coverage patterns similar to contigs assigned to Hemiptera (**Supplementary Figure 4**). Visualisation of coverage patterns and blastn hits along this contig with IGV v2.5.3 (Thorvaldsdóttir et al. 2012; Robson et al. 2012) showed a clear drop in PacBio coverage at 1,097,915 bp coinciding with a transition in blast hits from aphid sequences to Wolbachia sequences (**Supplementary Figure 7**). Given that the intersection between higher and lower coverage regions of the contig was only spanned by a single PacBio read, this contig was considered to be chimeric and was split at 1,097,915 bp and the Wolbachia section removed from the assembly. Additionally, contigs with less than 15x average MiSeq coverage and less than 10x average PacBio coverage were flagged as low coverage contaminants and removed. Finally, the mitochondrial (mt) genome was identified and removed based on alignment to the *M. persicae* mt genome (NCBI accession number KU877171.1) with nucmer v4.0.0.beta2 (Marçais et al. 2018), and patterns of coverage and GC content obtained from BlobTools.

### RNA-seq scaffolding

To further increase the contiguity of the updated *A. glyicnes* biotype 4 assembly, I performed RNA-seq based scaffolding of the filtered quickmerge assembly with P_RNA_scaffolder (Zhu et al. 2018). 5 Gb Paired-end RNA-seq data from Bansal *et. al.* (2014) was processed with Trim_galore! v0.4.5 to remove adapter sequences and trim bases with quality scores below 20, retaining paired reads that were at least 50 bp long after trimming. The trimmed RNA-seq data was aligned to the filtered quickmerge assembly with HISAT2 v2.0.5 “-k 3 --pen-noncansplice 1000000” (Kim et al. 2015) and the resulting SAM file passed to P_RNA_scaffolder which was run with default settings on the filtered quickmerge assembly. The scaffolded assembly was then sorted by scaffold length and scaffolds named numerically from longest to shortest to create a final release (A_gly_v2). A_gly_v2 was then checked for completeness and contamination with BUSCO and BlobTools respectively following the procedures outlined for A_gly_v1.

### RNA-seq informed gene prediction with BRAKER2

Prior to gene prediction, I soft-masked A_gly_v2 with known Insecta repeats from the RepeatMasker library using RepeatMasker v4.0.7 (Tarailo-Graovac and Chen 2009) with default settings. I then mapped the quality and adapter trimmed RNA-seq reads from Bansal *et. al.* (2014) (also used for genome scaffolding) to the soft-masked assembly with HISAT2 with the following parameters: “--max-intronlen 25000 --dta-cufflinks”. BRAKER2 v2.0.4 (Hoff et al. 2015, 2019) was then used to train AUGUSTUS (Stanke et al. 2008; Lomsadze et al. 2014) and predict protein coding genes, incorporating evidence from the RNA-seq alignments.

### Synteny analysis with A_gly_v1, A_gly_v2 and *D. melanogaster*

I investigated synteny between A_gly_v1 and A_gly_v2, and between A_gly_V2 and *Drosophila melanogaster* based on the identification of colinear blocks of genes with MCScanX v1.1 (Wang et al. 2012). For the *D. melanogaster* comparisons, I downloaded the R6.22 version of the genome assembly and gene predictions from FlyBase (Thurmond et al. 2018). In all cases, where multiple transcripts were annotated I used the longest transcript per gene as the representative transcript. For each comparison, I carried out an all vs. all blast search of annotated protein sequences using BLASTALL v2.2.22 with the options: “-p blastp - e 1e-10 -b 5 -v 5 -m8”. I then ran MCScanX, requiring at least three consecutive genes to call a colinear block (“-s 3”) and only searching for inter-species blocks (“-b 2”), all other settings were left as default. The MCScanX results were visualised with SynVisio (synvisio.github.io/#/) and the dual_synteny_plotter application from the MCScanX package.

### Annotation and analysis of *Osiris* genes

I used a combination of synteny, BLAST searches and phylogenetic analysis to annotate *Osiris* genes in A_gly_v1 and A_gly_v2. I extracted protein sequences of all annotated *Osiris* genes from the R6.22 *D. melanogaster* gene set and carried out blastp searches against A_gly_v1 and A_gly_v2 proteins with an e-value cut-off of 1 × 10^−5^. I then combined all identified *Osiris* proteins and aligned them with MUSCLE v3.8.31 (Edgar 2004) using default settings. Based on the MUSCLE alignment I carried out Maximum Likelihood (ML) phylogenetic analysis with FastTree v2.1.7 (Price et al. 2010) using default settings and visualised the resulting tree with FigTree (github.com/rambaut/figtree). A_gly_v1 and A_gly_v2 *Osiris* genes were named based on their closest relative to *D. melanogaster Osiris* genes in the ML tree. *A. glycines Osiris* genes without clear *D. melanogaster* ortholog were given and N(n) suffix. Syntenic relationships between A_gly_v1, A_gly_v2 and *D. melanogaster* with visualised using the SimpleSynteny web server (Veltri et al. 2016).

## Data availability

A summary of sequence data used in this manuscript is given in **Supplementary Table 3**. The final and intermediate genome assemblies (including *A. glycines* mitochondrial and endosymbiont genomes), and genome annotations generated for this study are available from Zenodo (doi:10.5281/zenodo.3453468).

## Acknowledgments

The described work was supported by a BBSRC Future Leader Fellowship (BB/R01227X/1) awarded to the author and CEPAMS grant 17.03.2 International Strategy Fund of the John Innes Centre. This research was supported in part by the NBI Computing Infrastructure for Science Group, which provides technical support and maintenance to the John Innes Centre’s high-performance computing cluster and storage systems. I thank Prof. Andy Michell and Dr. Jacob Wenger for providing access to sequence data from original *A. glycines* genome assembly project. I also thank Prof. Saskia Hogenhout for her support throughout the project and Prof. Cock Van Oosterhout for valuable comments on the manuscript.

