## Supplementary Figure for "Improved genome assembly and annotation of the soybean aphid (*Aphis glycines* Matsumura)"

**Supplementary Figures**

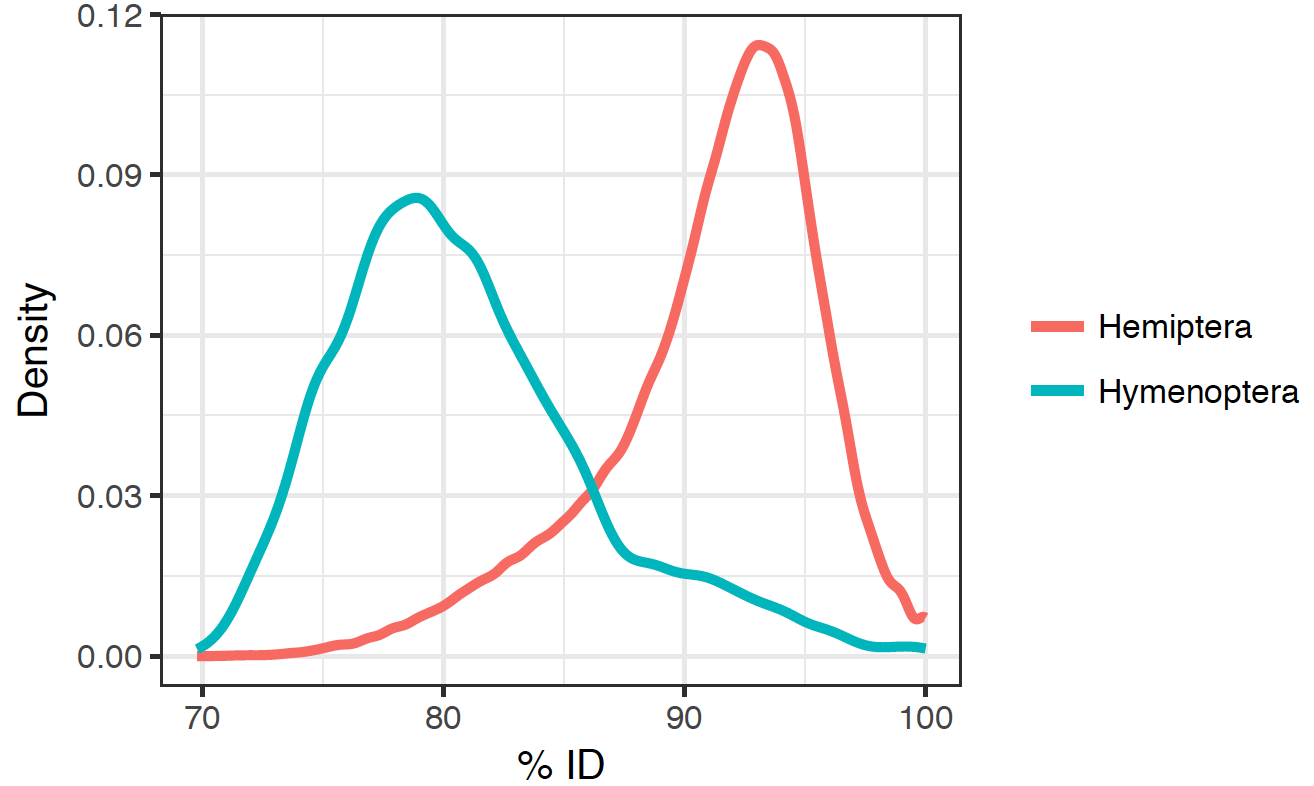

**Supplementary Figure 1:** The distribution of blast hit identities from sequences in the NCBI nucleotide database to scaffolds in A_gly_v1 assigned by BlobTools to either Hemipetera or Hymenoptera. See **Supplementary Table 1** for scaffold assignments.

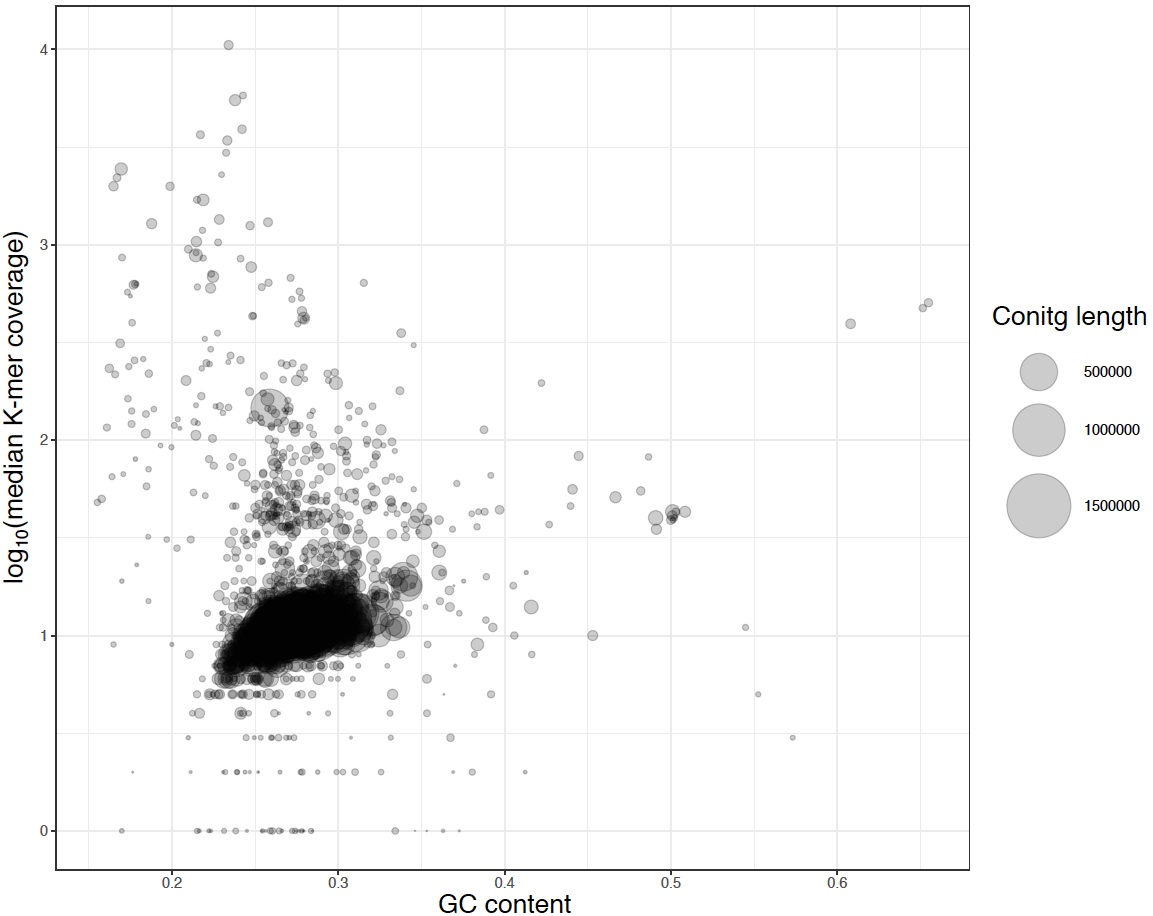

**Supplementary Figure 2:** GC content-coverage plot of the Canu assembly of *A. glycines* biotype 4 PacBio data. For each contig, the median K-mer coverage and GC content was calculated using KAT sect. K-mer coverage of each scaffold was calculated using all *A. glycines* biotype 4 Illumina Miseq libraries from Wenger et. al. (2017). Each circle in the plot represents a contig with the size of the circle scaled by contig length. Contigs form a primary “blob” around the expected GC content and coverage.

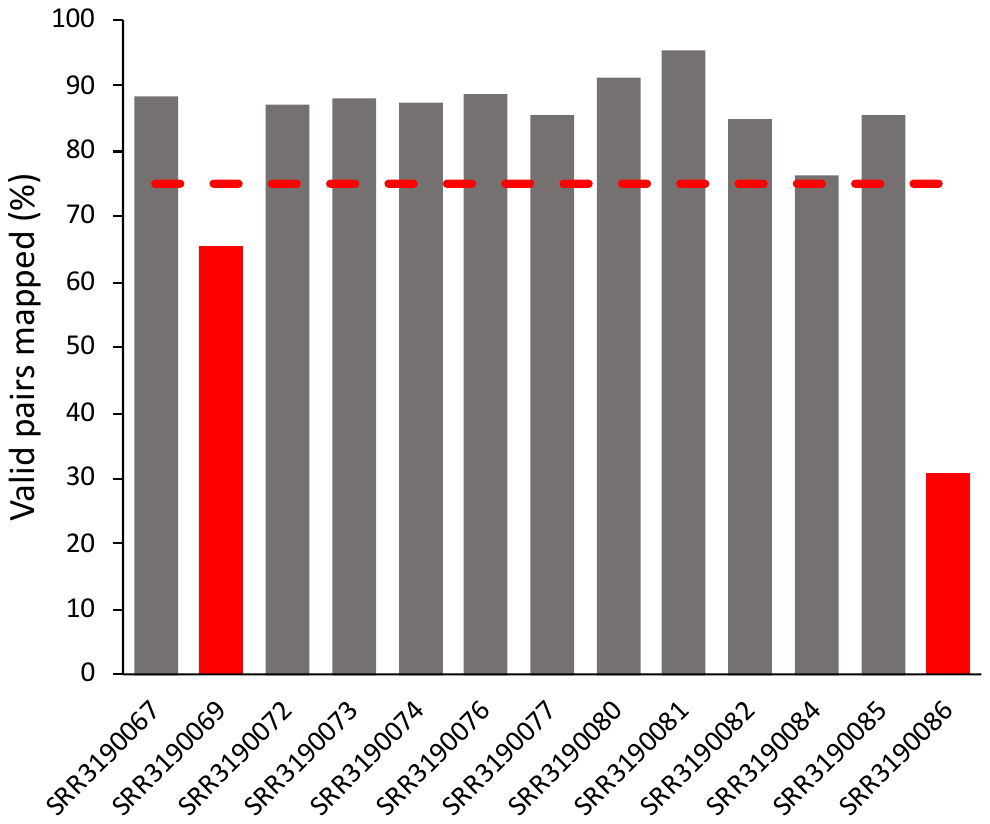

**Supplementary Figure 3:** Mapping results of *A. glycines* biotype 4 Illumina MiSeq libraries from Wenger et. al. to a Canu assembly of PacBio data derived from lab reared *A. glycines* bioptype 4 colony from the same study. Libraries with less than 75% mapping rates were not used for genome assembly.

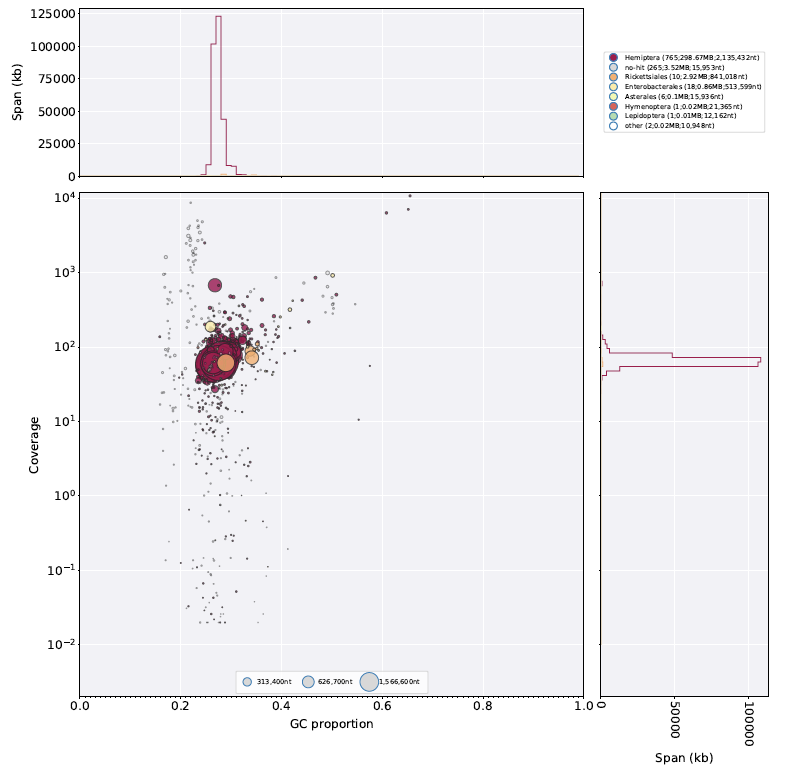

**Supplementary Figure 4:** BlobPlot of the unfiltered quickmerge assembly with taxonomy annotated at the Order level.

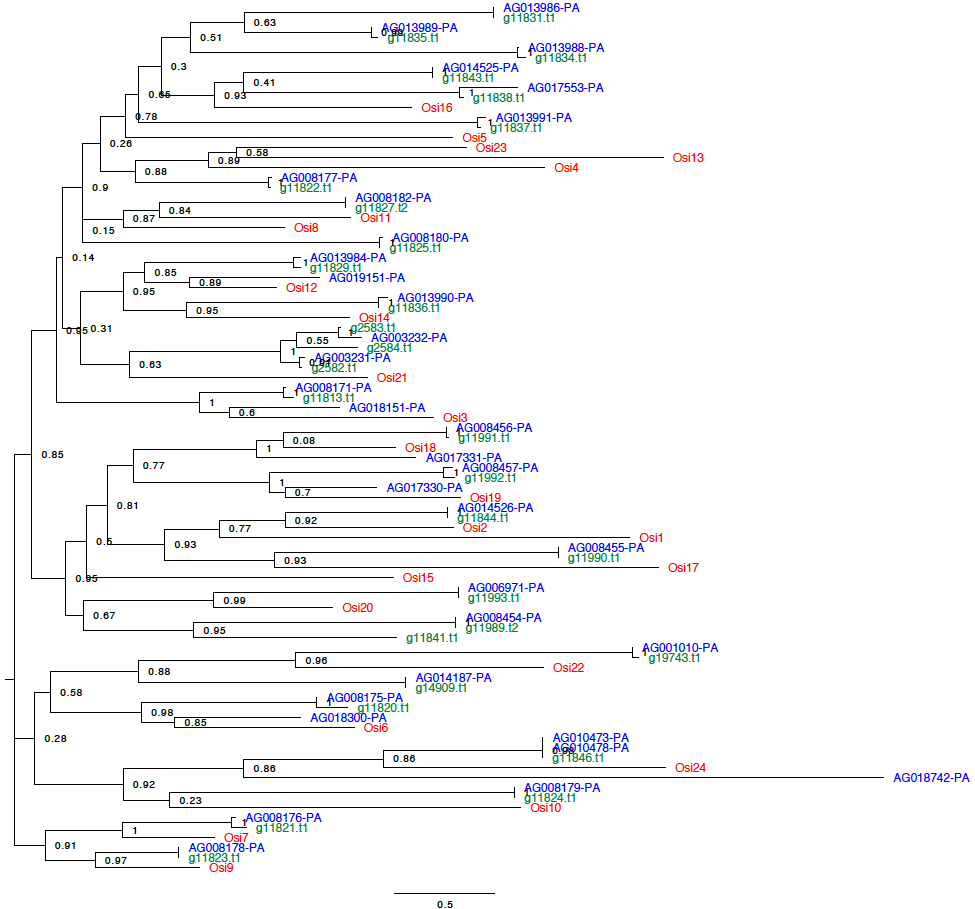

**Supplementary Figure 5:** Unrooted Maximum Likelihood phylogeny of *A. glycines* and *D. melanogaster* Osiris genes. Phylogeny estimated with FastTree based on a MUSCLE alignment of Osiris protein sequences (alignment available from Zenodo DOI: 10.5281/zenodo.3453468). Values at nodes indicate SH-like support values calculated by FastTree. *D. melanogaster* = red, A_gly_v1 = blue, A_gly_v2 = green. Osiris gene names assigned to A_gly_v1 and A_gly_v2 are summarised in **Supplementary Table 2**.

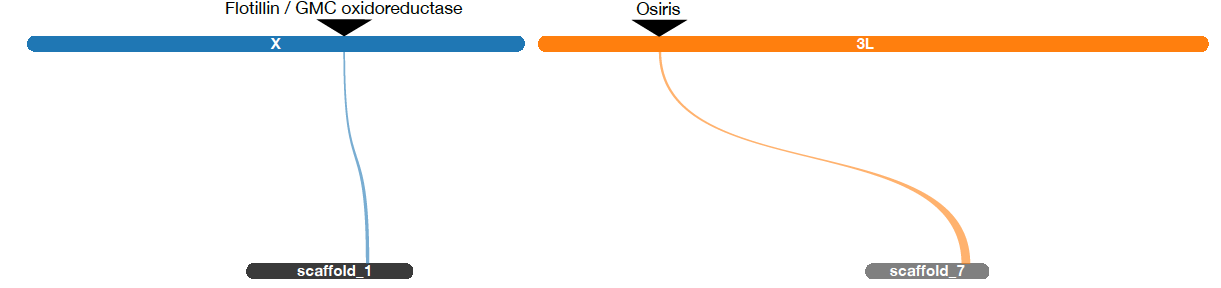

**Supplementary Figure 6:** SynVisio visualisation of syntenic blocks identified by MCScanX between *D. melanogaster* (top) and A_gly_v2 (bottom).

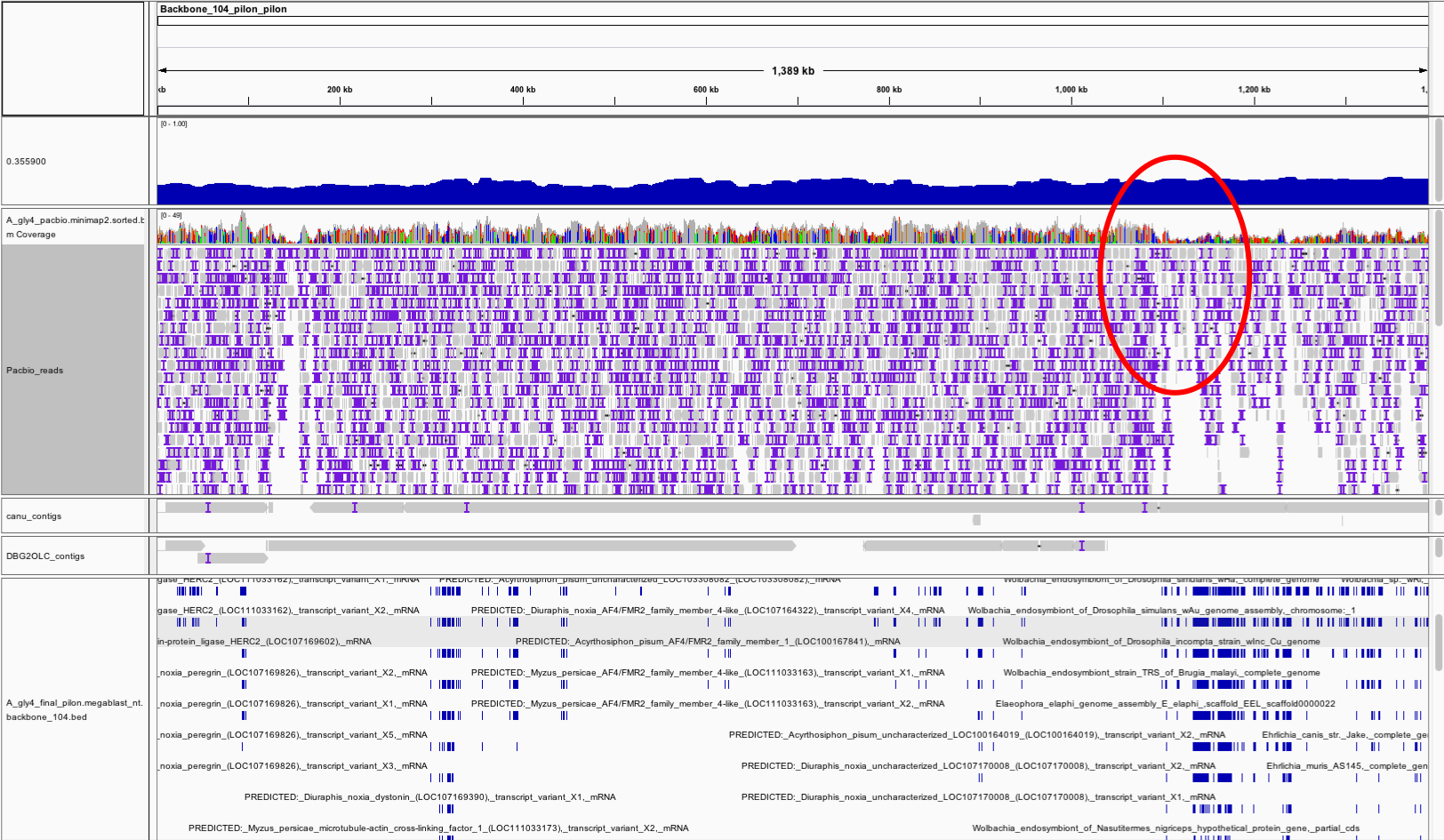

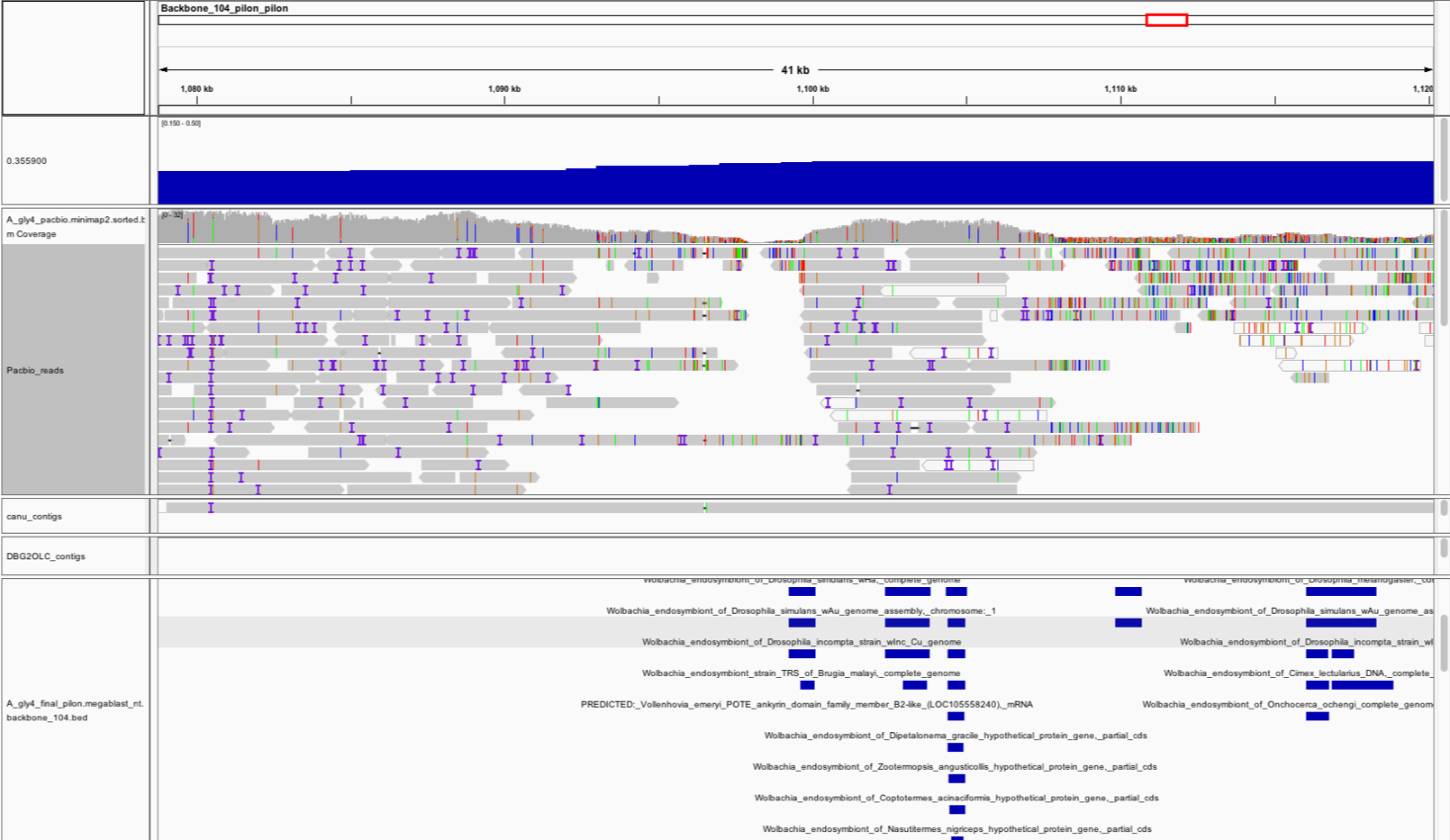

**Supplementary Figure 7:** IGV browser view of a putatively chimeric contig identified in the quickmerge assembly (backbone_104_pilon_pilon). Tracks from top to bottom: GC content along the contig, IGV coverage track of aligned PacBio reads, aligned PacBio reads, aligned contigs from the Canu assembly, aligned contigs from DBG2OLC assembly, NCBI nt database blastn hits. Red circle in top panel indicates region where coverage drops. Bottom panel is zoomed in view of the low coverage region.
